# NetWheels: A web application to create high quality peptide helical wheel and net projections

**DOI:** 10.1101/416347

**Authors:** A.R. Mól, M.S. Castro, W. Fontes

**Author notes:** Corresponding author. Full postal address: Laboratory of Protein Chemistry and Biochemistry, Department of Cell Biology, Institute of Biology, Bloco J, térreo University of Brasília, Brasília-DF, Brazil, 70910-900.

## Abstract

Helices are one of the most common secondary structures found in peptides and proteins. The wheel and net projections have been proposed to represent in two dimensions the tridimensional helical structures and facilitate the observation of their properties, especially in terms of residues polarity and intramolecular bonding. Nevertheless, there are few software options to create these projections. We have developed a web-based application that has several futures to create, customize and export these projections and is freely available at http://lbqp.unb.br/NetWheels.

## Introduction

Amino acid residues properties, such as polarity and acidity, play a fundamental role in defining peptides and proteins secondary structures, as well as their interactions with other substances. Alpha helices are one of the most common secondary structures found in these molecules. They arise from intramolecular interactions between the amino acid residues in peptides or proteins. In order to graphically represent these intramolecular interactions, different projections of the secondary structures of peptides have been created, most notably the wheel (Schiffer-Edmundson) and net projections (Schiffer and Edmundson 1967; Dunnill 1968).

Peptides helical wheel projections are drawn in such way that the observer sees the helical structure from the same axis which the helix grows. This is done by displaying each residue on the perimeter of a circle or spiral, with angles of 100° for every three residues, resulting in 18 resides until positions repeat, following the real conformation of alpha helixes. The wheel projection is mostly useful to visualize the helix regions by peptide properties, namely acidity/basicity and the ability to form hydrophilic (hydrogen) or hydrophobic bonds (Castro et al. 2006).

Net projections are used for the same structures as wheels, but provide a different perspective to the visualization of the helixes. They focus on displaying specific interactions between residues that are next to each other in terms of the helix central axis. The net projection is created by “cutting” the helix along its axis and displaying the residues in perpendicular to it. Intramolecular bonds between residues can be displayed, preferably using different patterns to differentiate the types of interactions.

Examples of these projections found in the literature vary a lot in terms of aesthetics (Dennison et al. 2005; Libério et al. 2011; Vicente et al. 2013; Mechkarska et al. 2013). Different color schemes have been used, as well as different shapes for polygons representing the residues as well as the lines representing their bonds. Nevertheless, these visual aspects are always used to represent the physicochemical properties of the peptide and highlight characteristics that can be of special interest for specific studies.

Different software have been developed to help researches draw and visualize helical wheel projections. Some of these are available as simple web applications and were not reported in journal publications (Armstrong and Zidovetzki; Everett; Kael Fischer; O’Neil and Grisham). Of those which were published, it was either as part of a larger bioinformatics package (Rice et al. 2000; Deléage et al. 2001; Gautier et al. 2008) or have been developed more than two decades ago using much simpler programming tools which were available at the time (Jones et al. 1992). A similar application to draw coiled coils has been published, but it does not cover the general helical wheel projections (Grigoryan and Keating 2008).

Even though these applications achieve the purpose of helping visualize the peptide characteristics, they lack the resources necessary to create high quality images ready for publications, especially when different aesthetics options are desired, such as customization of residues polygons, their colors or patterns, as well as size-related properties. While some focus on physicochemical properties of the helices, others are simple tools to visualize the disposition of the residues. We found only one report of a software capable of drawing net projections, however it has been published several years ago and isn’t available online (Jones et al. 1992).

Therefore, the aim of this web application is to provide an easy to use, open source yet powerful tool to create peptide wheel and net projections, which can be highly customizable and exported in high quality graphics formats.

## Material and Methods

The projections are drawn using the powerful plotting engine of the R programming language and environment (R Core Team 2015), which allow for a detailed customization of every figure aspect necessary to get the desired output. The web application, which works as a guided user interface (GUI) for the R plotting functions was created using the shiny package extended with the shinyjs package (Attali 2015; Chang et al. 2015). These packages make the web application development almost uniquely done using the R language, which greatly simplifies its development and maintenance.

## Results and Discussion

The web application is accessible and its source code available freely at http://lbqp.unb.br/NetWheels. Its use is very straightforward, with options grouped in tabs (Fig. 1) that are used to set parameters for both wheel and net projections.

**Fig. 1.**
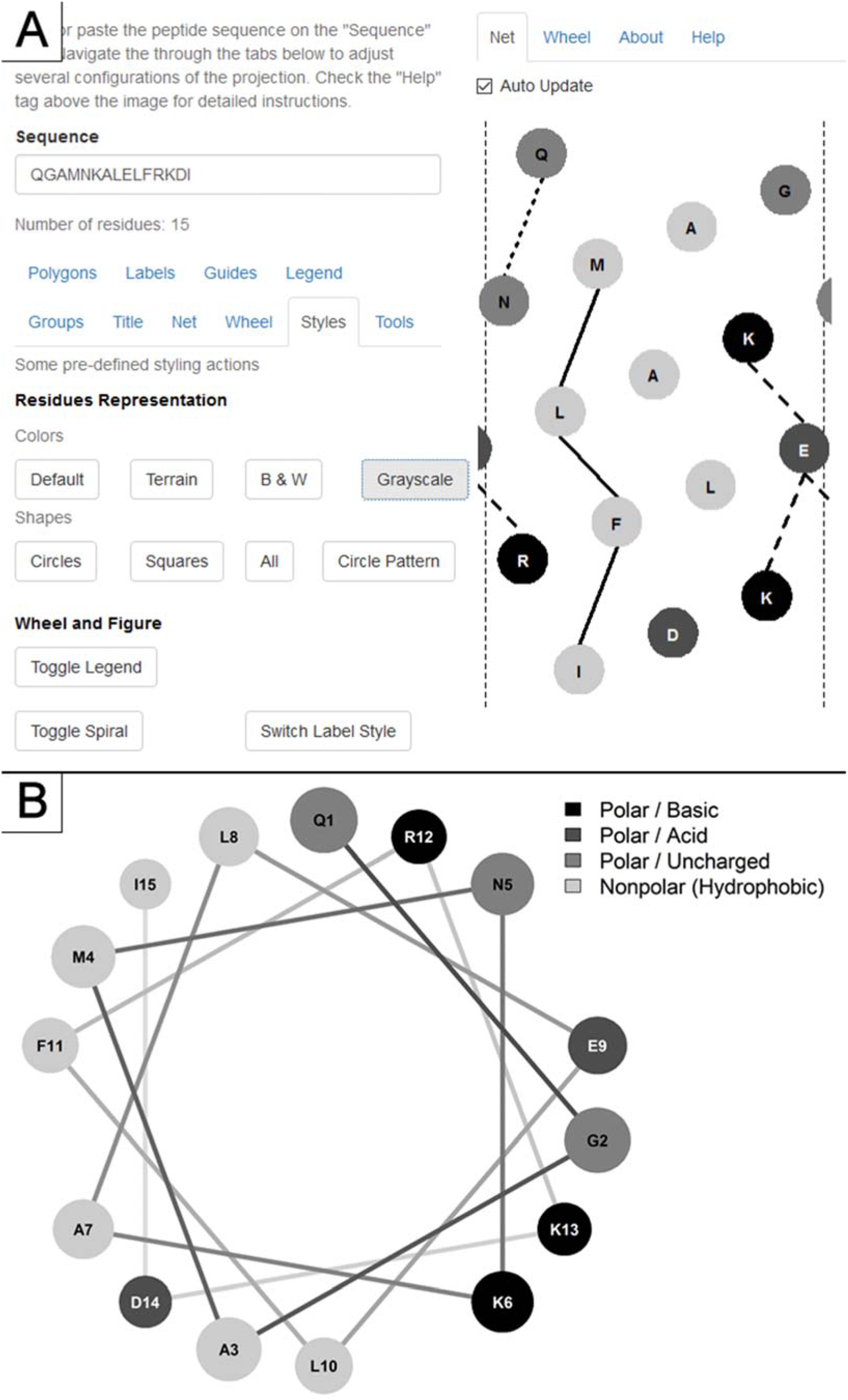
Screenshot of the application showing the tabs used to set the parameters and a net projection (A) and the wheel projection (B) for the same peptide.

The amino acid residues are classified in four groups: basic, acid, uncharged polar and nonpolar. There is also the possibility of specifying an unknown residue group. Each group can have its aesthetics properties adjusted individually, such as polygon type, fill (colors and patterns) and borders, and also labeling properties regarding position, font-size and amino acid code style.

A set of pre-defined styles are available in order to facilitate the use for those without strict customization necessity. Parameters related to the nature of α-helices, namely the 18 residues period and 3.6 residues per turn, can also be changed in order to enable the projection of 3_10_ and π helices. Reproducibility is guaranteed by the option to export all the parameters used to create a given projection, making it easy to load them back in and adjust the projections if necessary, without having to manually reset them. The projections can be exported to different image formats and resolutions, which enables their direct use in documents and presentations without loss of quality.

We believe that the NetWheels application can be of great value for those looking to design high-quality customized projections

## Acknowledgements

The authors acknowledge CNPq, FINEP, CAPES, FAP-DF and FUB-UnB for financial support.

## Conflict of interest and involvement of human or animal subjects

The authors declare that there is no conflict of interest regarding the publication of this manuscript. The authors declare that there were no human or animal subjects in the present work as it is only theoretical and computational.

